# Novel Hendra virus variant detected by sentinel surveillance of Australian horses

**DOI:** 10.1101/2021.07.16.452724

**Authors:** Edward J. Annand, Bethany A. Horsburgh, Kai Xu, Peter A. Reid, Ben Poole, Maximillian C. de Kantzow, Nicole Brown, Alison Tweedie, Michelle Michie, John D. Grewar, Anne E. Jackson, Nagendrakumar B. Singanallur, Karren M. Plain, Karan Kim, Mary Tachedjian, Brenda van der Heide, Sandra Crameri, David T. Williams, Cristy Secombe, Eric D. Laing, Spencer Sterling, Lianying Yan, Louise Jackson, Cheryl Jones, Raina K. Plowright, Alison J. Peel, Andrew C. Breed, Ibrahim Diallo, Navneet K. Dhand, Philip N. Britton, Christopher C. Broder, Ina Smith, John-Sebastian Eden

## Abstract

A novel Hendra virus (HeV) variant, not detected by routine testing, was identified and isolated from a Queensland horse that suffered acute, fatal disease consistent with HeV infection. Whole genome sequencing and phylogenetic analysis demonstrated the variant to have ~83% nucleotide identity to the prototype HeV strain. An updated RT-qPCR assay was designed for routine HeV surveillance. *In silico* and *in vitro* comparison of the receptor-binding protein with prototypic HeV showed that the human monoclonal antibody m102.4 used for post-exposure prophylaxis, as well as the current equine vaccine, should be effective against this variant. Genetic similarity of this virus to sequences detected from grey-headed flying-foxes suggests the variant circulates at-least in this species. Studies determining infection kinetics, pathogenicity, reservoir-species associations, viral–host co-evolution and spillover dynamics for this virus are urgently needed. Surveillance and biosecurity practices should be updated to appreciate HeV spillover risk across all regions frequented by flying foxes.

## Introduction

Highly pathogenic zoonotic Hendra virus (HeV) and Nipah virus (NiV) are the prototypic members of the genus *Henipavirus*, family *Paramyxoviridae*, with natural reservoirs in pteropodid flying-foxes (1). These viruses exhibit wide mammalian host tropism, causing severe acute respiratory and encephalitic disease mediated by endothelial vasculitis, with high fatality and chronic encephalitis among survivors (2–4). As of March 2021, 63 natural spillovers of HeV had been recognized in Australian horses resulting in 105 horse deaths (5, 6), with four of seven confirmed human cases having been fatal (7). NiV has caused zoonotic outbreaks in South Asia and Southeast Asia with very high case fatality rates (70-91%) resulting in more than 700 human deaths (8–10). In response to the zoonotic disease threat posed by henipaviruses to humans and domestic animals, vaccines and post-exposure prophylaxis (PEP) have been developed (11). This includes a licensed subunit vaccine (Equivac^®^ HeV) used in Australian horses since 2012, which is based on a soluble version of the HeV receptor-binding protein (RBP; attachment G glycoprotein) (HeV-sG) (12). A human monoclonal antibody (mAb) m102.4 has been administered as emergency PEP in 16 human cases and has demonstrated safety, tolerability and intended immunogenicity in phase 1 trials (13). Combinations of cross-reactive humanized antibodies against the fusion (F) protein and the RBP have also been described for clinical development as PEP (14–16), and a human vaccine using HeV-sG is now in phase 1 clinical trials (17).

Horses are the predominant species known to be infected with HeV by natural spillover from flying-foxes, with two canine (18) and all known human infections having resulted from close contact with infected horses. HeV transmission from *Pteropus* spp. (flying-foxes) to horses is thought to occur primarily via contaminated urine (19). The spatial distribution of previously detected spillovers to horses, as well as molecular HeV testing of flying-fox urine, suggested that transmission was predominantly from black flying-foxes (BFF, *P. alecto*) and spectacled flying-foxes (SFF, *P. conspicillatus*) (19, 20). However, serological testing has detected antibodies to HeV or related henipaviruses amongst all four Australian flying-fox species, with the reported seroprevalence ranging between ~20% and ~65% (20–23). Notably, seroprevalence of IgG targeting the HeV RBP in *P. poliocephalus*, the grey-headed flying-fox (GHFF), has been reported as 43% in South Australia and Victoria (22) and 60% (169/284) in south-east Queensland (21).

Australia hosts more-than 1 million horses. The grazing behavior of horses, their large respiratory tidal volume, and an extensive highly-vascularized upper respiratory epithelium all contribute to their vulnerability for HeV spillover (23). Detection of spillover to horses relies on the clinical recognition of consistent disease by attending veterinarians, appropriate sampling, and laboratory submission for priority state laboratory testing (24). Passive surveillance via suspect disease testing is affected by a strong regional bias for areas where HeV has previously been detected, and where domestic horse populations overlap with the distribution of BFFs, ranging from eastern coastal Queensland to northern New South Wales (25). Testing for HeV is less commonly performed on horses suffering similar disease further south in Australia due to a perception that the likelihood of disease occurrence is lower in regions without this flying-fox species (26).

Despite high rates of HeV testing in horses with consistent disease across regions of established risk (>1000 tested annually), less than 1% are found to be positive (25, 27).

Routine testing for equine HeV infection as part of priority disease investigation is specific for the matrix (M) gene (28). Additional nucleoprotein (N) gene specific testing (29) is limited to HeV-positive samples that undergo confirmatory testing (30) or in the minority of suspected equine HeV cases submitted directly to the national reference laboratory from states where spillover was considered less likely (<7% nationally) (25) and state testing was unavailable. Importantly, this means that most horse-disease cases found to be negative for HeV are not investigated further. This is despite evidence for other viruses circulating with potential spillover risk to Australian horses, including novel related bat-borne paramyxoviruses (27, 31–35). This is consistent with animal health surveillance globally, which often prioritizes exclusion testing targeting pathogens of established significance rather than open-ended diagnostic approaches that are inherently more challenging to implement and interpret.

Using a multidisciplinary and interagency approach combining clinical–syndromic analysis, molecular and serological testing, we explored the hypothesis that some severe viral-like diseases of horses consistent with HeV, but negative when tested, result from undetected novel-paramyxovirus spill-over from flying-foxes, potentially posing similar zoonotic risk. Here we report the identification of a previously unrecognized variant of HeV (HeV-var) that resulted in severe neurological and respiratory disease in a horse, clinically indistinguishable from prototypic HeV infection.

## Materials and Methods

### Study cohort

A biobank of diagnostic specimens was developed from horses that underwent HeV testing in Queensland between 2015 and 2018 but were negative for HeV by reverse transcriptase quantitative PCR (RT-qPCR) (28). Clinical, epidemiological and sample-related data were recorded including vaccination status and perceived exposure to flying-foxes as variably reported by submitting veterinarians. All samples were archived at −80°C. We applied a decision algorithm based on pathogenic-basis and syndromic analysis of clinical disease to categorize each case by likelihood of having an infectious viral cause (Appendix Table). Samples (EDTA blood, serum, nasal swab, rectal swab) from cases assigned the highest likelihood of having infectious cause (priority categories 1 & 2) were plated for serological screening and high-throughput nucleic acid extraction using the MagMAX^™^ mirVANA and CORE pathogen kits (ThermoFisher, Australia).

### Pan-paramyxovirus RT-PCR screening

cDNA was prepared from extracted RNA using Invitrogen SuperScript IV VILO mastermix with ezDNase (ThermoFisher, Australia). A nested reverse transcriptase-PCR (RT-PCR) assay targeting the paramyxovirus L protein gene was adapted using primers developed by Tong et al (36) and the Qiagen AllTaq PCR Core kit (QIAGEN, Australia). Amplicons corresponding to the expected size (584 bp) were identified by gel electrophoresis before purification with AMPure XP (Beckman Coulter, Australia). To capture any weak detections, pools were also prepared by equal volume mixing of all PCR products across plated rows. Next-generation sequencing was performed using an Illumina iSeq with the Nextera XT DNA library preparation kit (Illumina, Australia). For analysis, reads were assembled with MEGAHIT (37) before identification by comparison to NCBI GenBank with BLAST (38).

### HeV-var whole-genome sequencing

Samples positive by paramyxovirus RT-PCR for the novel HeV-var were subjected to meta-transcriptomic sequencing to determine the complete genome sequence and identify any co-infecting agents. RNA was reverse-transcribed with Invitrogen SSIV VILO mastermix (ThermoFisher, Australia) and FastSelect reagent (QIAGEN, Australia). Second-strand synthesis was performed with Sequenase 2.0 before DNA library preparation with Nextera XT and unique dual indexes. Sequencing was performed on an Illumina NextSeq (Illumina, Australia) generating 100 M paired reads (75 bp) per library.

### Assembly and comparative genomic and phylogenetic analyses

For genome assembly, RNA sequencing reads were trimmed and mapped to a horse reference genome (GenBank GCA_002863925.1) using STAR aligner to remove host sequences. The non-host reads were *de novo* assembled with MEGAHIT (37) and compared with the NCBI GenBank nucleotide and protein databases using blastn and blastx (38). The putative virus contig was extracted and reads were remapped to this draft genome using bbmap v37.98 (https://sourceforge.net/projects/bbmap) to examine sequence coverage and identify misaligned reads. The majority consensus sequence was extracted, aligned and annotated by reference to the prototype HeV strain using Geneious Prime v2021.1.1, and submitted to GenBank (accession number MZ318101).

For classification, the paramyxovirus polymerase (L) protein sequence was aligned according to International Committee on Taxonomy of Viruses (ICTV) guidelines (39). We prepared alignments of partial nucleocapsid (N) and phosphoprotein (P) nucleotide sequences with known HeV strains from the GenBank database. Phylogenies were prepared using a maximum likelihood approach in MEGA X (https://www.megasoftware.net/) according to the best-fit substitution model and 500 bootstrap replicates.

### RT-qPCR assay development

An existing RT-qPCR assay targeting the HeV M gene (28) was adapted to target the novel HeV-var. The duplex assay used the Applied Biosystems AgPath-ID One-Step RT-PCR kit (ThermoFisher, Australia) and distinguishes prototype and variant HeV strains. Briefly, 4 μL RNA was combined with 10 μL 2× RT-PCR Buffer, 0.8 μL 25× RT-PCR Enzyme Mix, 2 μL nuclease-free water and 3.2 μL primer/probe mix (0.6 μL each primer, 0.3 μL each probe from 10 μM stock; Table 1). The reaction was performed as follows: 10 min at 50°C for cDNA synthesis, 10 min at 95°C for RT inactivation, and 50 cycles of 95°C for 15s and 60°C for 30s with FAM and HEX channels captured at the end of each cycle. As positive control, gene fragments were synthesized encoding a T7 promoter upstream of the partial M gene for both prototypic and variant HeV (Supplementary Figure 1). RNA transcripts were expressed using the NEB HiScribe T7 High Yield RNA Synthesis kit (New England Biolabs, Australia).

**Table 1.**
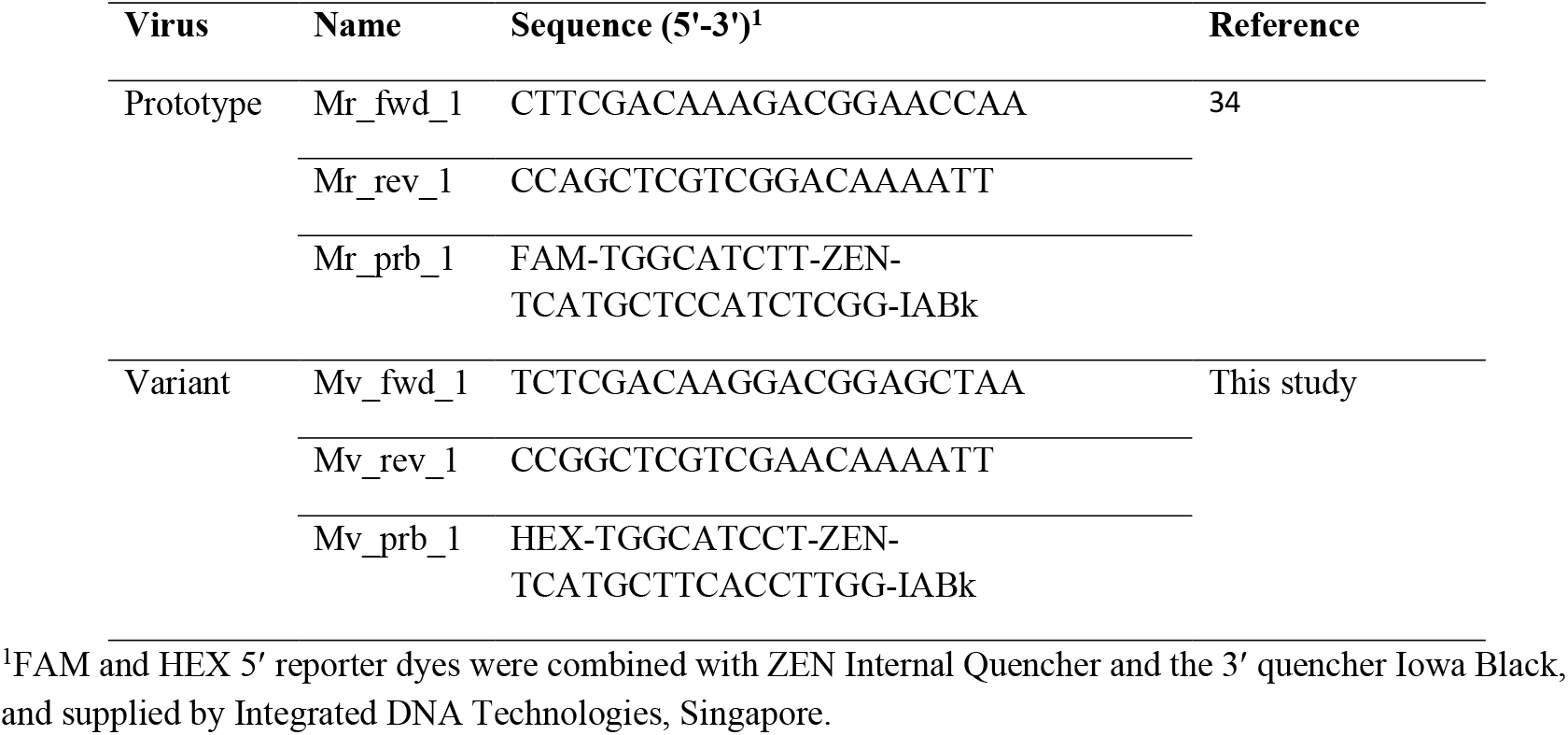
Oligonucleotides used for new HeV duplex RT-qPCR targeting matrix gene

**Figure 1.**
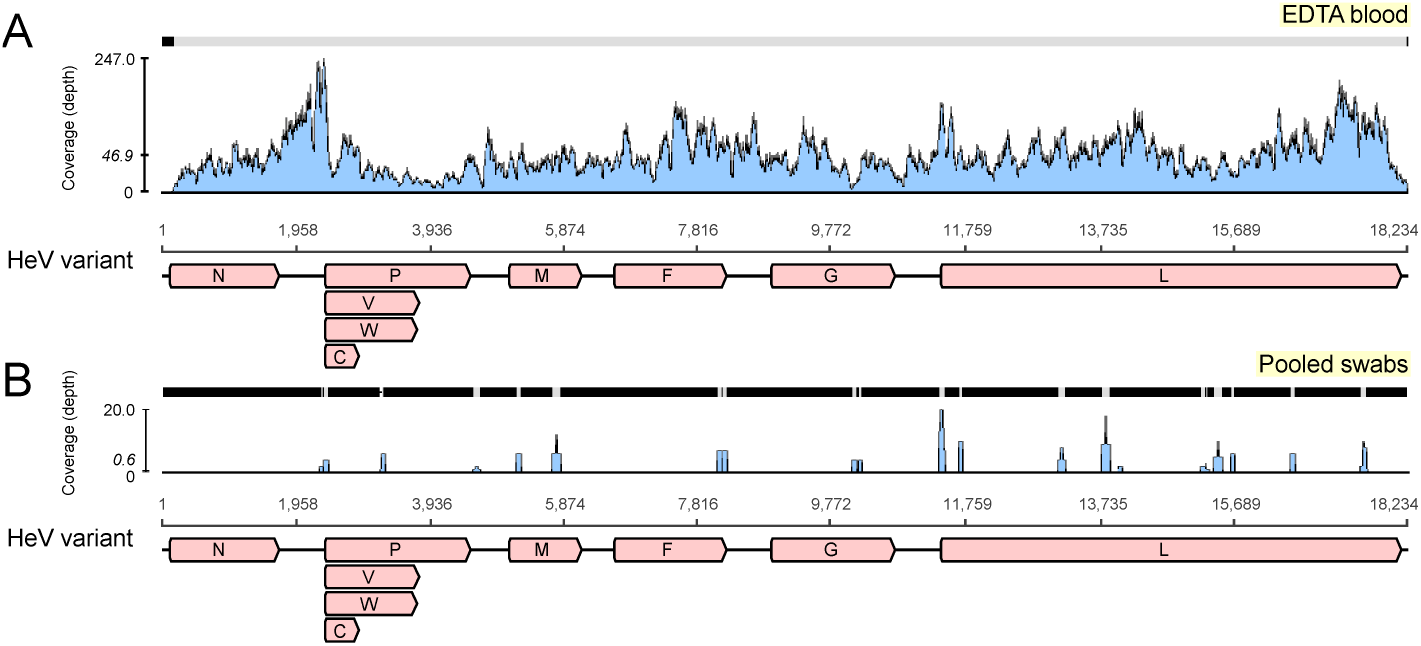
Sequence coverage of novel Hendra virus (HeV) variant. The RNA sequencing reads were mapped to the novel HeV variant genome to examine coverage across the genome and depth for the EDTA blood and pooled swab samples in panels A and B, respectively. The x-axis shows the genome position with genes annotated and the y-axis shows the sequence read coverage (depth). The italicized coverage value indicates the mean coverage depth for that sample.

### Virus isolation, confirmation and neutralization studies

Isolations were attempted in Vero cells (ATCC CCL-81) and primary kidney cells derived from *P. alecto* (40), and confirmed by CPE formation, RT-qPCR, RNA sequencing, electron microscopy and viral neutralization studies using mAb m102.4 against HeV and isolated variant (Supplementary methods).

### Serological analysis

Serological analysis was performed using multiplex microsphere immunoassays with a Luminex MAGPIX^™^ system (ThermoFisher, Australia). Initial screening for IgG antibodies was undertaken using an extensive panel of bacterial (*Leptospira*, *Brucella*) and viral antigens (paramyxovirus, filovirus, coronavirus, flavivirus, alphavirus) coupled to MagPlex beads (Bio-Rad, Australia) for multiplex screening. Briefly, blood or serum, diluted 1:100, was added to the beads, with binding detected following the addition of a combination of biotinylated-Protein-G and -A and streptavidin–R–phycoerythrin. The median fluorescence intensity (MFI) was read on the MagPix (Luminex) targeting 100 beads per antigen with a Bayesian latent class model used to assess test performance and determine appropriate cut-offs for positive test classification (32). An immunoglobulin (Ig)M assay was also applied in which biotinylated anti-equine IgM antibodies were used in place of biotinylated Protein A and G.

### In silico *analysis of the RBP homology and mAb binding*

The translated protein sequence of the HeV-var RBP sequence was compared with the X-ray crystal structures of the HeV RBP protein structure bound to mAb m102.4 (41) and to ephrin-B2 using SWISS-MODEL (https://swissmodel.expasy.org/) to assess the ability of m102.4 to neutralize this variant and further-establish the likelihood of antibodies produced by immunization with the HeV vaccine being protective against this variant.

## Results

### Case report

In September 2015, a 12-year-old Arabian gelding in south-east Queensland presented with severe disease consistent with HeV infection. The horse had always resided on the affected property and disease onset was acute with rapid deterioration over 24 h. Clinical assessment determined depressed (obtunded) demeanor, injected/ congested gingival mucous membranes (darkened red/ purple change with darker periapical line and prolonged capillary refill time), tachycardia (75 beats/min), tachypnoea (60 breaths/min), normal rectal temperature (38.0°C), muscle fasciculations, head pressing and collapse.

HeV infection was suspected by the attending veterinarian, who had previously managed a confirmed case, based on consistent clinical disease and perception of plausible flying-fox exposure. A near-by roost is known to host BFFs, GHFFs and little red flying-fox species with numbers varying seasonally and annually (42). Nasal, oral and rectal swab samples were obtained postmortem and combined in 50mL of sterile saline, and blood was collected in an EDTA tube. The moribund condition justified euthanased on humane grounds.

Pooled swabs and blood samples were submitted to the Queensland Biosecurity Sciences Laboratory for priority HeV testing. Testing for HeV by RT-qPCR targeting the M gene did not detect viral RNA and IgG antibodies against the HeV RBP were not detected by ELISA (28, 43).

### Identification of novel HeV-var

Given the case’s assigned high-likelihood of zoonotic infectious cause (Supp table 1), both the EDTA blood and pooled swab samples were screened by pan-paramyxovirus RT-PCR (36). This identified the partial polymerase sequence of a novel paramyxovirus, most closely related to HeV (≈ 11% nucleotide difference). Deep sequencing of blood RNA generated the near-full length genome of a novel HeV (mean coverage of 46.9x) (Figure 1A). The virus was less abundant in the pooled swab sample, with a mean coverage depth of 0.6X reads spanning only 9.9% of the genome (Figure 1B). Importantly, no other viruses were present in either sample, and other microbial reads assembled were from common microflora including *Staphylococcus aureus*, *Aeromonas*, *Veillonella*, *Pseudarthrobacter*, *Streptococcus*, *Acinetobacter* and *Psychrobacter* species.

### Confirmation of HeV infection

A comparison of the primer and probe sequences used for routine diagnostic assay (28, 29) revealed multiple mismatches in the binding sites, explaining the failure of routine surveillance to detect this variant (Figure 2). An RT-qPCR assay was designed to detect both prototype and variant HeV strains in duplex (Table 1, Supplementary Figures 1, 2), which amplified the templates of each virus with similarly high efficiency (>94%) and sensitivity - capable of detecting <100 copies of target RNA (Supplementary Figure 2). EDTA blood and pooled swabs samples were quantified, confirming RNA sequencing, with higher abundance in the EDTA blood than in the pooled swabs (quantification cycle (Cq) 26.87 versus 30.67 respectively). No further cases were identified by re-screening the priority cohort (864 samples representing 350 Queensland cases) using this novel assay.

**Figure 2.**
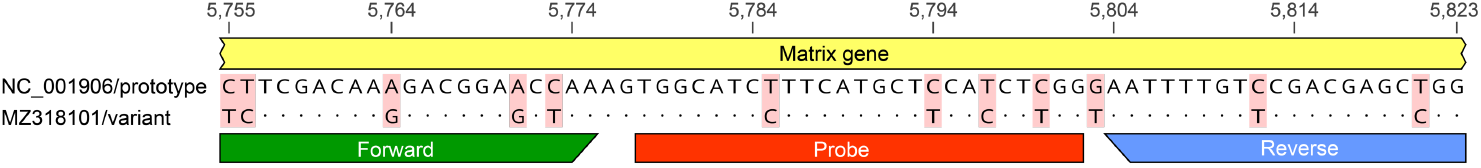
Genomic variation in the Hendra virus (HeV) matrix gene assay primer/probe binding sites. The genomic region targeted by the commonly used HeV matrix gene RT-qPCR assay (Smith et al *J Virol Methods* 2001;98:33–40) was aligned and compared for the prototype and variant HeV strains. The genomic positions relative to the prototype strain (NCBI GenBank accession NC_001906) are shown at the top. Primers (forward and reverse) and probe binding sites are indicated by the colored bars. Mismatches between the sequenced have been highlighted with red shading.

Virus isolation from the EDTA blood sample of the case animal was successful in Vero cells. Electron microscopy of infected cells revealed cytoplasmic inclusion bodies (nucleocapsid aggregations) (Figure 3A), and enveloped viral-particle budding (Figure 3B), consistent with HeV (Figure 3 A-D) (44).

**Figure 3.**
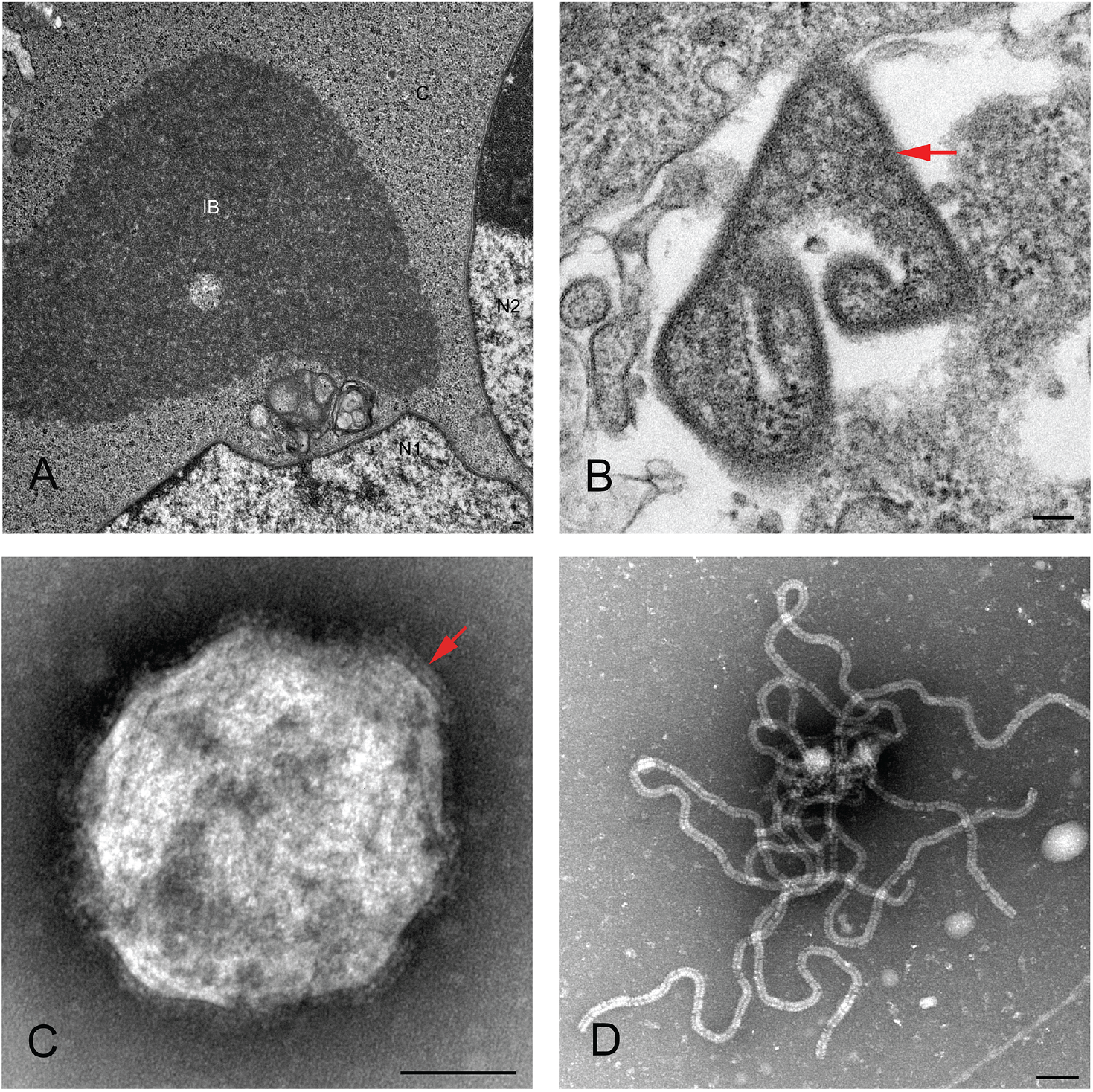
Transmission electron micrographs of Vero cells inoculated with the EDTA blood sample. Panel A and B are thin section, Panel C and D are negative contrast analyses. Panel A shows an inclusion body (IB) within the cytoplasm (C) of a multinucleated (N1 and N2) syncytial cell. The non-membrane bound IB consists of hollow nucleocapsids. Panel B, red arrow indicates virion with egress occurring at the plasma membrane. Panel C, red arrow indicates a double fringed envelope of the virion. Panel D shows strands of ribonucleic protein (RNP) characteristic of the family *Paramyxoviridae*. Scale bars represent 100nm.

Blood was tested for IgM and IgG antibodies against a panel of 33 antigens representative of bacterial and viral zoonoses (32, 45), including RBPs of paramyxoviruses: HeV, NiV, Cedar henipavirus (CedV), Mojiang henipavirus (MojV, Ghana bat henipavirus (GhV), Menangle pararubulavirus, Grove and Yeppoon viruses. No significant reactions were observed for this case in either the IgG or IgM assays, indicating a lack of detectable antibodies consistent with acute viremia.

### Genomic analysis of novel HeV-var

Phylogenetic analyses of the novel HeV-var was performed with other known paramyxoviruses (Figure 4A–C). Comparison of the nucleotide similarity of the novel HeV-var to the HeV prototype strain (GenBank NC_001906) revealed an 83.5% pairwise identity across the genome (Figure 4D). Importantly, the L protein phylogeny revealed that the branch lengths of prototype and variant HeV to their common node did not exceed 0.03 substitutions/site (Figure 4A, B), thus according to ICTV criteria the viruses are considered of the same species (39). However, the HeV-var is clearly well outside known HeV diversity (Figure 4C).

**Figure 4.**
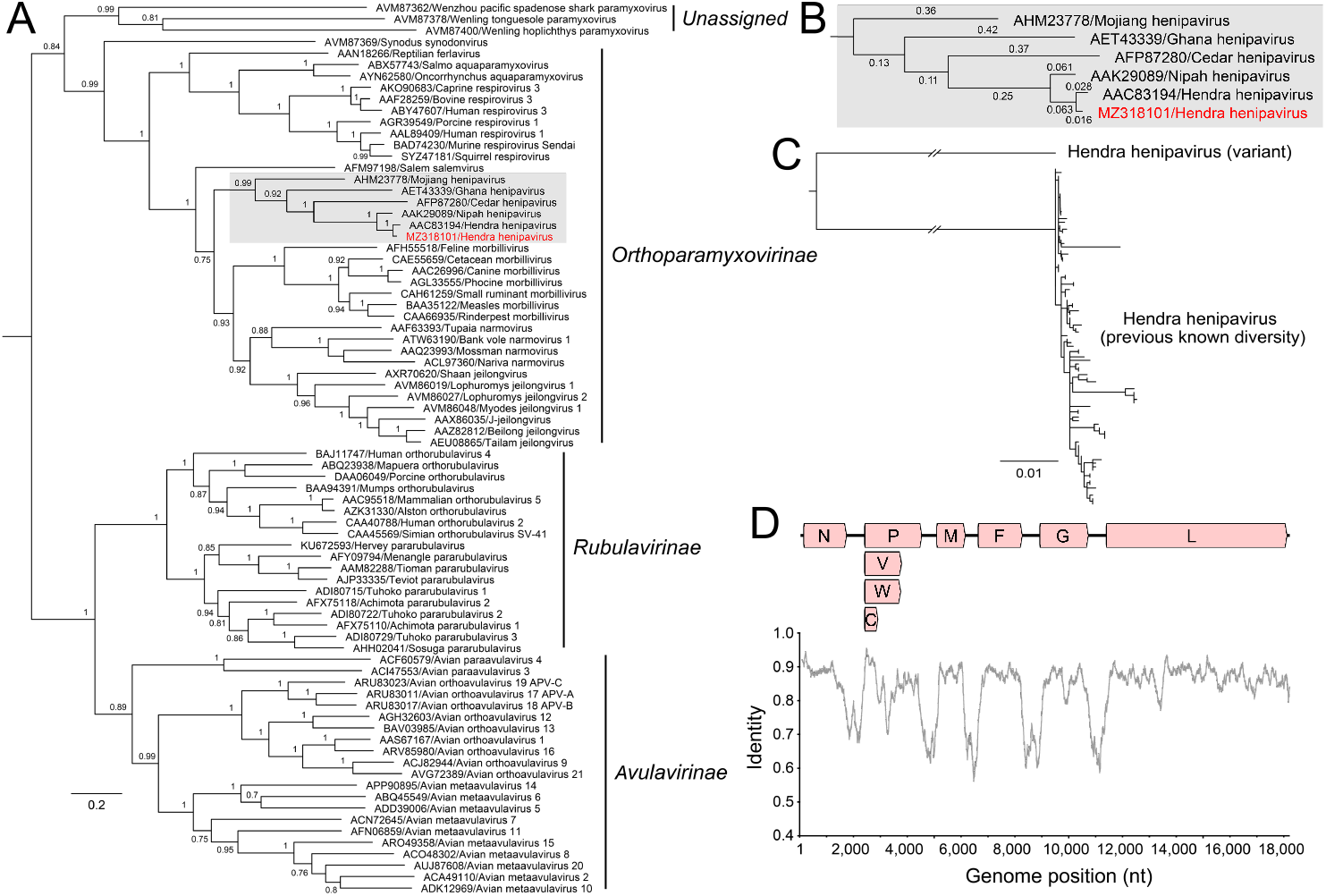
Phylogenomics of novel Hendra virus (HeV) variant. Panel A shows a maximum-likelihood phylogeny of paramyxoviruses using complete L protein sequences with henipaviruses highlighted in grey. The novel HeV (colored red) groups with the prototypic HeV. Bootstrap support values as proportions of 500 replicates are shown at nodes with values less than 0.7 hidden. Panel B shows the branch lengths for the henipavirus clade, showing that the branch leading back to the common ancestor of all known HeVs and this novel virus does not exceed 0.03; thus they are considered variants of the same species. For the phylogenies, the scale bars indicate the number of substitutions per site. Panel C shows a maximum-likelihood phylogeny of partial N and P where deep branch lengths have been collapsed for visualization only to demonstrate that the variant is well outside the known diversity of HeV. Panel D shows nucleotide genomic similarity of the variant compared with the prototypic HeV strain.

Following this finding, comparison with a partial novel Henipavirus M gene sequence derived from a GHFF from South Australia in 2013 (46) revealed 99% similarity to this HeV-var. This, along with subsequent further flying-fox detections (47), suggests that this HeV-var represents a previous undescribed lineage, with reservoir-host infection across at-least the range of this flying-fox species.

### Analysis of the RBP

Genomic sequence showed greatest variability in the non-coding regions with mean pairwise genome identity higher (86.9%) across coding regions (Figure 4D). At the protein level, this HeV-var shared between 82.3% and 95.7% amino acid identity (mean 92.5%) to the HeV prototype (Table 2). Notably, the HeV-var RBP shared 92.7% amino acid identity with prototypic HeV. Modeling of the novel HeV-var RBP structure based on the translated protein sequence using the previously published X-ray crystal structure of the prototypic HeV RBP (40) revealed that the epitopes for binding of ephrin-B2 receptor, as well of mAb m102.4, should remain functionally unchanged due to consistency between key residues (Figure 5). Indeed, mAb m102.4 neutralization assays conducted in parallel with HeV and HeV-var revealed equivalent neutralization potency of m102.4 (2.3μg/mL of m102.4 neutralized 30TCID_50_ of HeV-var and 4.6 μg /mL of m102.4 neutralized 300TCID_50_ of HeV).

**Table 2.** Novel HeV variant protein length and similarity to prototype strain (NCBI GenBank accession NC_001906)

| Protein | Length (AA) | Similarity (%) |
| --- | --- | --- |
| Nucleoprotein | 532 | 96.6 |
| Phosphoprotein | 707 | 82.3 |
| Matrix | 352 | 95.7 |
| Fusion | 546 | 95.4 |
| Glycoprotein | 603 | 92.5 |
| Large | 2,244 | 95.7 |

**Figure 5.**
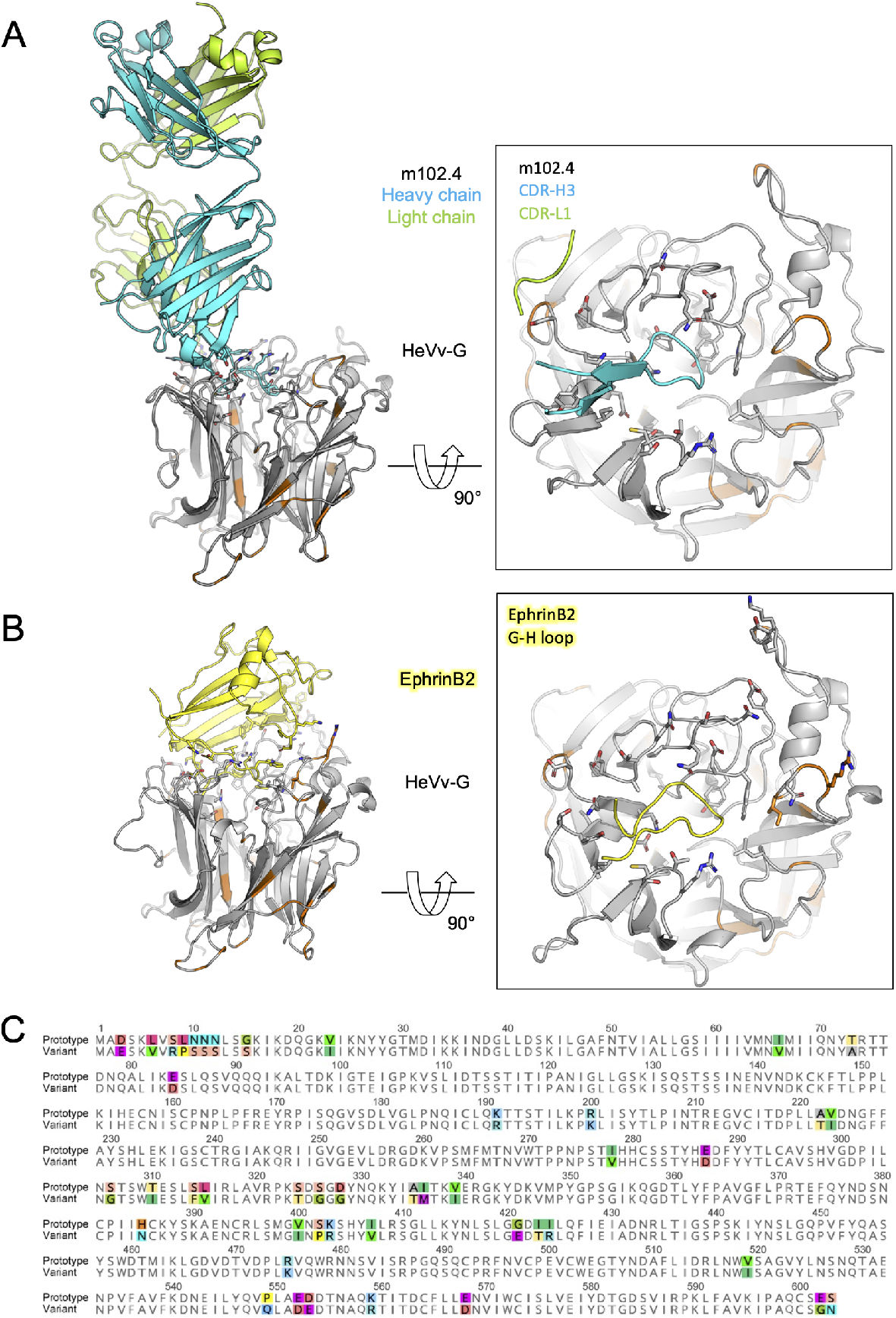
*In silico* modelling of Hendra virus (HeV) variant G protein (receptor-binding protein) with ephrin-B2 and the human monoclonal antibody m102.4. The translated protein sequence encoded by the HeV variant G gene was modelled using the known protein structure of the prototype virus bound to the human monoclonal antibody m102.4(A), and the receptor ephrinB2 (in yellow) (B). On the left of both panel A and B show the side views of the interactions between the HeV G protein and the two binding partners, highlighting key binding residues (as sticks) and the variant positions (colored in orange) relative to the m102.4 (heavy chain in cyan and light chain in green) and ephrinB2 (in yellow). The insets of both panel A and B show zoomed top views at the HeV G and m102.4/ephrinB2 binding interface, highlighting specific interactions by the complementarity-determining regions of the mAb and G-H loop of the receptor ephrinB2. These data show that variable positions do not occur at critical epitopes at the HeV G and m102.4 binding interface and have very minor impact to the receptor ephrinB2 binding. Panel (C) shows an alignment of the prototypic and variant HeV strain G proteins with variable positions highlighted in color.

## Discussion

We describe a significant virus discovery resulting from diagnostic investigations as part of an innovative, targeted, syndromic surveillance extending-from routine priority disease investigation. Based on the ICTV criteria (39), the virus identified is a novel genotypic variant of HeV, rather than a new henipavirus species. The novel HeV-var missed detection by routine diagnostic testing due to genomic divergence yet demonstrated equivalent infection and disease consistent with its phenotypic similarity to HeV. The findings highlight potential for improved emerging pathogen detection via ‘One Health’ interagency and multidisciplinary syndromic infectious disease research and prompt recognition of the importance of targeted emerging infectious disease (EID) sentinel surveillance. We have described a new assay for routine human and animal health laboratory diagnosis and surveillance of this virus.

Comparison of the translated amino-acid sequence of this HeV-var and prototypic HeV RBP *in silico* revealed the ephrin-B2 entry receptor binding site and that of mAb m102.4 to be unchanged. Similarly, m102.4 neutralization was confirmed *in vitro* with HeV-var as for HeV. As such, it is expected that current PEP utilizing mAb m102.4 will remain effective against this HeV-var. It should be emphasized, that the HeV RBP shares only 79% amino acid identity to NiV RBP, yet the HeV-sG subunit vaccine provides 100% protection against lethal challenge with both HeV and NiV in animal models (11). Both the higher similarity between the HeV-var and HeV RBP (92.5% amino acid identity) and structural consistency of critical epitopes mentioned above, suggest that current vaccination utilizing the HEV RBP will elicit similarly protective antibodies against this HeV-var. Current serological assays based on the HeV RBP are not expected to distinguish between exposure to the variants.

The 99% similarity between this HeV-var and a partial M-gene sequence detected in a GHFF from Adelaide in 2013, highlights that: a greater diversity of HeV circulates among Australian flying-fox species than has been previously recognized; and that this novel variant likely circulates as a relatively consistent sub-lineage, at-least across the range of this flying-fox species. Additional identifications in GHFF and LRFF from regions previously lacking in molecular HeV detections have subsequently further supported this understanding (47).

These findings prompt urgent reassessment of HeV spillover risk for horses and handlers living in southern NSW, Victoria and South Australia, where previously the risk has been considered substantially lower than regions within the distribution of BFFs. Our findings indicate a need to update molecular assays, increase disease surveillance testing in horses, and screening of flying-foxes for this HeV-var in these areas. This may further resolve the previously reported anomaly of high seropositivity within these species despite low HeV detection (20–22).

Despite relatively high genetic divergence, the phenotypic similarity of this variant to the previously known prototypic HeV, combined with the observed consistency of disease in the horse, suggests they are of equivalent pathogenicity and spillover potential. Further characterization of HeV genomic diversity and any host-species associations will increase our understanding of transmission dynamics as well as viral–host co-evolution features such as possible co-divergence or founder effects. Indeed, BFFs have rapidly expanded their range southward in response to habitat clearing and other anthropogenic influences (48) - increasing their overlap with GHFFs. Climatic change and habitat loss are altering the extent and nature of interspecies interactions. Sampling of multiple species across time and space is needed to understand how this HeV-var circulates within and among flying-fox species. Clearly, biosecurity practices should be updated to include appreciation of spillover risk in all regions frequented by flying-foxes, regardless of species.

Passive disease surveillance and biosecurity risk management for emerging diseases relies on recognition and management of suspect disease cases by clinical veterinarians in private practice who play crucial roles relevant to animal and human health (24). Surveillance for HeV and ABLV in Australian horses is inherently challenged by their high zoonotic consequence yet infrequent sporadic and rare incidence respectively. Prioritization of this case in our research testing pathway was based on clear description of disease consistent with HeV by the attending veterinarian. Veterinarians are challenged in performing such disease recognition by obligations to simultaneously: serve the animal and animal owner; manage biosecurity risk; and meet *Workplace Health and Safety Act and Biosecurity Act* requirements (24, 49, 50). This detection highlights the potential for improved EID surveillance via parallel serology and molecular testing pathways constructed to suit available sample-types and target diseases of highest clinical, species and geographic relevance when guided by systematic interpretation of clinical and field observations made as part of existing submission and biosecurity protocols (50, 51). The example serves a proof of concept for other significant disease contexts, highlighting benefit in integration of multidiscipline research approaches with routine biosecurity operations.

Acknowledging the limitations of this single case, that lacked tissue for histopathology and immunohistochemistry, it is nonetheless appropriate that this HeV-var be considered equally pathogenic to previously-known HeV based on coherent and consistent available evidence. Specifically: the observed clinical signs of disease and pathology; evidence of viraemia; the phylogenetic analysis indicating that the variant belongs to the HeV species; and the modeling of the interactions of the functional RBP domain to the virus entry ephrin-B2 receptor support understanding of consistent pathogenicity relative to the prototypic HeV. Moreover, this case fits the case definition for HeV infection in Australia’s AUSVETPLAN, which is that an animal tests positive to HeV using one or more of PCR, virus isolation or immunohistochemistry (51).

Updated PCR diagnostics suitable for routine priority detection of this HeV-var have been developed and are now in use in many of Australia’s animal and human health laboratories. These findings demonstrate the limitation of exclusion-based testing for emerging zoonoses and a gap in our understanding of how frequently detection of known zoonoses across a broad range of systems are missed because of the diagnostic tools used. Further investigations to determine the prevalence of HeV-var circulation among and excretion from all Australian flying-fox species should be prioritized. The risk of zoonotic HeV disease in horses and in-contact humans should be interpreted across all regions frequented by flying-foxes (regardless of species), particularly in areas previously considered to be low risk for HeV spillover.

## Author Biographies

Dr Annand is an equine veterinarian epidemiologist in private practice and a Research Associate at the University of Sydney School of Veterinary Science, Marie Bashir Institute of Infectious Diseases and Biosecurity. His research interests include One Health infectious disease surveillance and risk management.

Dr Horsburgh is an early-career researcher at the Westmead Institute for Medical Research. She is interested in using single-copy assays and high-throughput sequencing to characterize viral genomes and understand their effect on human health.

## Acknowledgments

The authors are grateful to: The staff of Biosecurity Sciences Laboratory, Brisbane, QDAF, Queensland for their work in processing of samples and submission information for this case and others in the comparative cohort. From CSIRO Australian Centre of Disease Preparedness (ACDP), Ms Jennifer Barr for processing of samples, Ms Andrea Certoma, and Mrs Mel Hargreaves for technical assistance in the isolation of the virus and Dr Jianning Wang for assistance with sequence comparisons. Mr David Bath, Dr Robyn Martin, Dr William Wong and others from the Department of Agriculture Water and the Environment involved with supporting this project as part of the Biosecurity Innovation Program. A/Prof. Jenny-Ann Toribio for PhD supervision (EA). Allan and Lyn Davies and their family DALARA foundation, for their philanthropic financial support through the founding years of this research - without which this knowledge gap might not have been closed for a great time longer. Veterinarians and horsepersons will be better informed in their management of this deadly zoonotic disease because of your support. We especially thank the veterinarians, owners and carers who manage horse health and that of the public in relation to HeV.

## Disclaimers

CCB is a United States federal employee and inventor on US and foreign patents pertaining to soluble forms of Hendra virus and Nipah virus G glycoproteins; and monoclonal antibodies against Hendra and Nipah viruses, whose assignees are The United States of America as represented by the Department of Health and Human Services (Washington, DC) and Henry M. Jackson Foundation for the Advancement of Military Medicine Inc. (Bethesda, MD). The remaining authors declare no competing interests. The opinions or assertions contained herein are the private ones of the author(s) and are not to be construed as official or reflecting the views of any of the Australian or international affiliated government or research agencies.

## Supplementary Methods

### Viral isolation

Positive case samples for the novel HeV were sent to the Australian Centre for Disease Preparedness (ACDP), a World Organization for Animal Health Reference Laboratory for Hendra and Nipah virus diseases, in line with established national arrangements for confirmatory testing of notifiable disease of animals. Virus isolation was attempted in Vero cells (ATCC CCL-81) and primary kidney cells derived from *P. alecto* (PaKi; (39) on whole blood and pooled nasal, oral and rectal swab samples. Vero cells were grown at 37°C in EMEM (ThermoFisher Scientific) containing 10% fetal calf serum (FCS; ThermoFisher Scientific), supplemented with 1% v/v L-glutamine, 10 mM HEPES, 0.25% v/v penicillin–streptomycin and 0.5% v/v amphotericin B (Sigma-Aldrich). PaKi cells were cultured in DMEM/F-12 media (ThermoFisher Scientific) with 5% FCS and supplemented as above.

For virus isolation, washed monolayers of cells were inoculated with 500 μL of whole blood diluted 1:5 in culture media or 500 μL of pooled swab sample prefiltered (0.45-μm cellulose acetate) to remove bacteria and any residual solid particles. Inoculum was removed after 45 min and cell monolayers were washed with phosphate-buffered saline, then overlaid with culture media containing 1% (v/v) FCS. Flasks were incubated at 37°C for 6–7 days and regularly monitored for cytopathic effect by light microscopy. Cells were then frozen, thawed and the cell suspension clarified by centrifugation (1000*g* at 4°C). Supernatant (500 μL) was then passaged onto fresh cell monolayers. A maximum of three passages per sample were performed on each cell line. Final pass samples were tested by RT-qPCR to detect the presence of replicating HeV genome.

### Electron Microscopy

For negative contrast EM: The clarified supernatant from Vero cell cultures, infected with HeV-var, were inactivated with 4% formaldehyde overnight. After adsorption of the inactivated supernatant onto formvar/carbon coated Cu400 grids, the preparation was then stained with Nano-W (Nano-probes) for 1min. For thin section EM: The pelleted cells were fixed with modified Karnovsky’s fixative (4% formaldehyde and 2.5% glutaraldehyde in 0.1M Sorensen’s phosphate buffer) at 4°C overnight. The pellet was rinsed in analogous buffer, fixed with 1% osmium tetroxide for 1 hr and dehydrated with a graded ethanol series prior to being embedded in Spurr’s resin (ProSciTech) according to the manufacturer’s instructions. A Leica UC7 microtome was used to produce ultrathin sections, which were then stained in saturated uranyl acetate in 50% ethanol followed by lead citrate. Grids were examined and images acquired using a JEOL JEM-1400 transmission electron microscope at 120V.

### Neutralization assay

Serial dilutions of the mAb m102.4 and Hendra virus (isolate Hendra virus/Australia/Horse/2008/Redlands) or the HeV-var (Hendra-var/Australia/Horse/2015/Gympie) diluted to contain 100 TCID_50_/well were incubated for 45–60 min at 37°C in a 96 well plate. A suspension of Vero cells was added to every well at a concentration of 4 × 10^5^ cells per ml. Positive and negative serum controls and virus-only controls were included. Plates were incubated in a humid atmosphere containing 5% CO_2_. Cells were examined after 3 days under an inverted microscope for cytopathic effect.

---

**Appendix Table.** Infectious disease prioritization categories (with examples) used in this study to identify Hendra-negative equine disease cases with highest likelihood of similar undiagnosed viral cause from larger cohort for further investigation

| Infectious disease priority | Description | Example |
| --- | --- | --- |
| *Category 1*<br>Highest infectious disease suspect | Case features ‘pyrexia’ or ‘abnormal mucous membranes AND one or more other clinical signs related to infectious disease OR the presence of either change AND ‘epidemiological observation indicative of infectious cause’ based on temporal and/or spatial relationship to similar disease cases | Pyrexia with tachycardia and acute onset respiratory consolidation and/or secretions. Pyrexia and neurological symptoms. Pyrexia and ‘injected/congested’ mucous membranes. ‘Congested/injected mucous membranes’ with acute severe respiratory dysfunction. Clustering of similar cases on same or neighboring properties |
| *Category 2*<br>High infectious disease suspect | Pyrexia OR other clinical signs associated with infectious disease of interest | Acute onset abnormal respiratory secretions. Fever of unknown origin. Colic with the presence of neurological symptoms |
| *Category 3*<br>Moderate infectious disease suspect | Clinical signs may be associated equally with infectious and non-infectious causes | Colic with the presence of dehydration and mucous membrane changes. Ataxia with the absence of pyrexia or known trauma |
| *Category 4*<br>Low infectious disease suspect | Non-infectious etiologies more common or most likely on differential diagnosis list, but infectious cause still possible | Ataxia following known traumatic event. Traumatic wounds following unusual behavioral event. Acute lethargy following chronic non-infectious disease condition |
| *Category 5*<br>No infectious disease suspect | No clinical signs of illness or no infectious cause considered likely | Traumatic wounds in the absence of underlying disease. Screening in unvaccinated horses to manage biosecurity risk prior to invasive procedures addressing non-infectious disease such as is a common requirement for dentistry or admission to equine hospitals in Australia |
| *Category 6*<br>Confirmed infectious disease | Other infectious disease confirmed via diagnostic testing | A case submitted for HeV testing, found negative and then testing positive for alternative known infectious disease such as ABLV, WNV, EHV or RRV* |

586 \*ABLV, Australian bat lyssavirus; EHV, Equine herpes virus; HeV, Hendra virus; RRV, Ross River Virus; WNV, West Nile virus

**Supplementary Figure 1.**
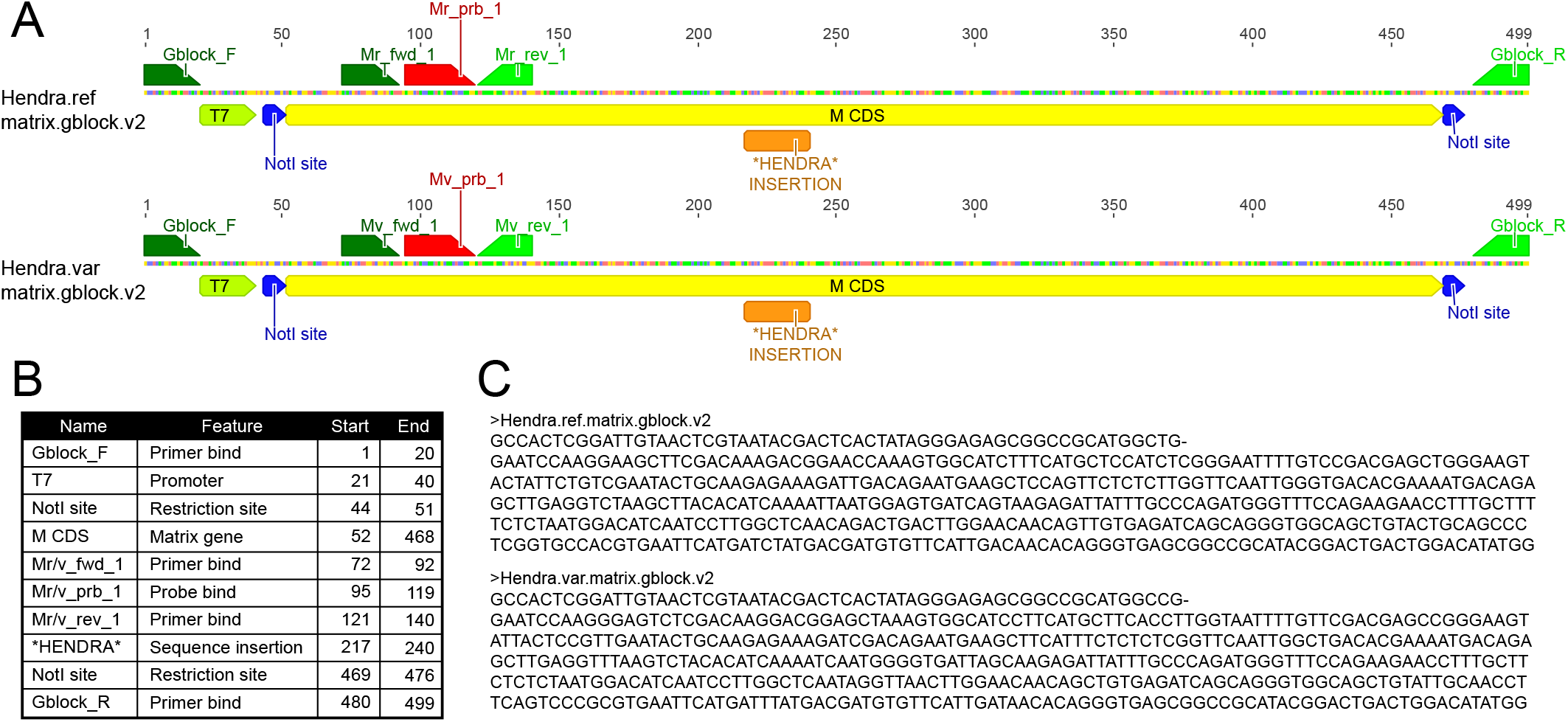
Synthetic genes used for positive controls in HeV RT-qPCR assay. Panel A shows genomic maps of the controls. The standards can be used as DNA or following T7 RNA transcription used as RNA standard for detection of HeV prototype (top) and variant (bottom) strain with M gene assay. The design and length for each are same except the equivalent M coding region has been swapped. The M coding regions have been flanked by regions from PCR amplification (Gblock F & R), T7 transcription, and cloning (NotI sites). Positions with feature keys and fasta format sequences have been providided in panels B & C, respectively.

**Supplementary Figure 2.**
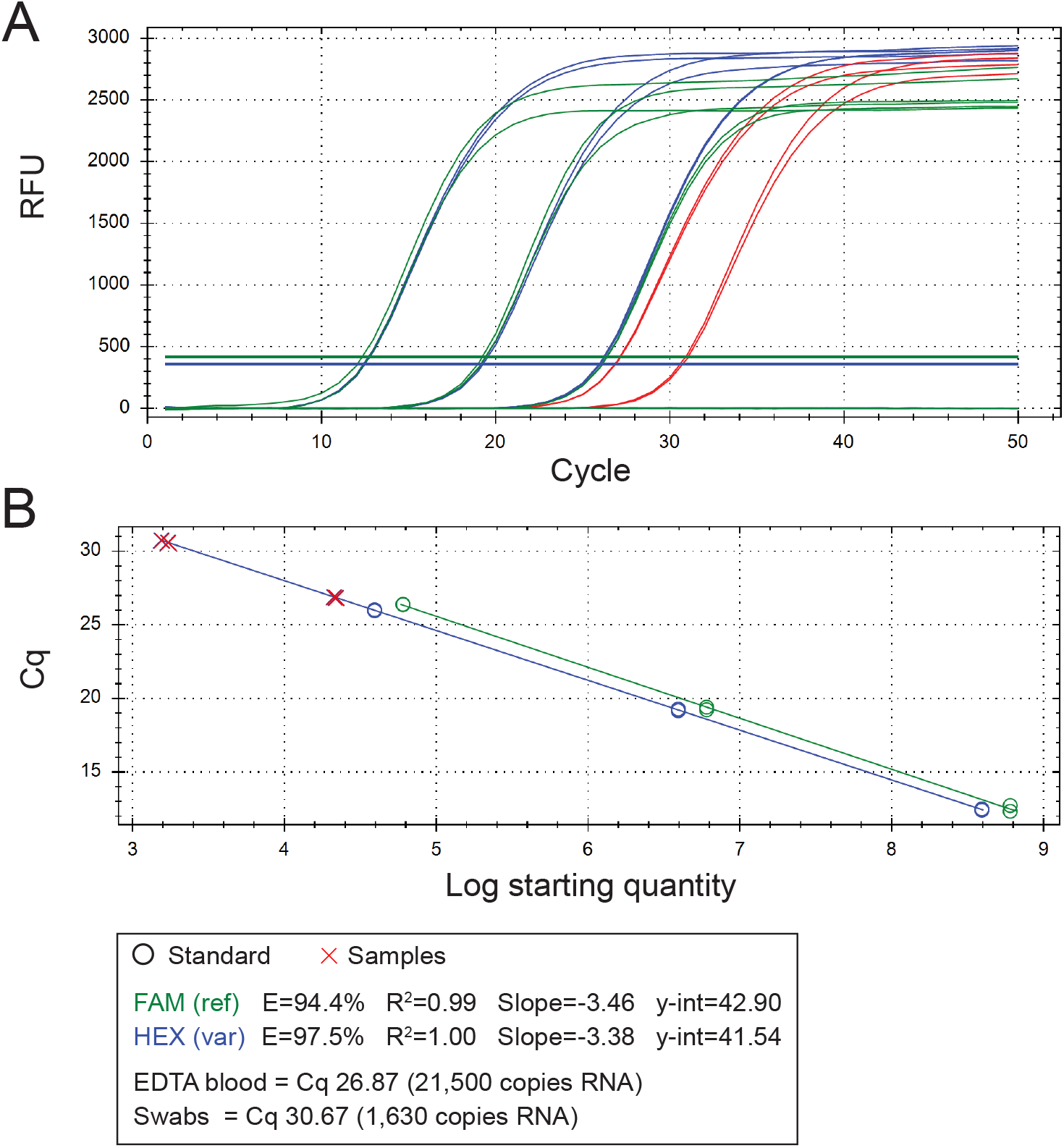
RT-qPCR performance and results for clinical samples. Panel A shows an amplification plot for the duplex RT-qPCR assay reporting both FAM & HEX channels (both HeV strains) with the synthetic RNA controls - green (FAM - reference strain) & blue (HEX - variant strain). The red colored traces are detections in the HEX (variant) channel for clinical samples of case p15_13388 EDTA blood and swab samples, respectively. Panel B shows the standard curve plot for the duplex RT-qPCR assay reporting both FAM & HEX channels (both HeV strains) with the synthetic RNA controls - green (FAM - reference strain) & blue (HEX - variant strain). Shapes, colors and results have been summarised the panel at the bottom.

## References

1. Eaton BT, Broder CC, Middleton D, Wang L-F. Hendra and Nipah viruses: different and dangerous. Nature Reviews Microbiology. 2006 Jan;4(1):23–35.

2. Selvey LA, Wells RM, McCormack JG, Ansford AJ, Murray K, Rogers RJ et al. Infection of humans and horses by a newly described morbillivirus. Medical Journal of Australia 1995;162:642–645.

3. Wong KT, Tan CT. Clinical and pathological manifestations of human henipavirus infection. Current Topics in Microbiology and Immunology. 2012;359:95–104.

4. Playford EG, McCall B, Smith G, Slinko V, Allen G, Smith I, et al. Human Hendra virus encephalitis associated with equine outbreak, Australia, 2008. Emerging Infectious Diseases. 2010 Feb;16(2):219–23.

5. Murray K, Rogers R, Selvey L, Selleck P, Hyatt A, Gould A, et al. A novel morbillivirus pneumonia of horses and its transmission to humans. Emerging Infectious Diseases. 1995 Mar;1(1):31–3.

6. Business Queensland. Summary of Hendra virus incidents in horses. 2015 [cited 2020 Feb 14]. Available from: https://www.business.qld.gov.au/industries/service-industries-professionals/service-industries/veterinary-surgeons/guidelines-hendra/incident-summary

7. New South Wales Health. Summary of human cases of Hendra virus infection - Control guidelines. [cited 2021 Apr 23]. Available from: https://www.health.nsw.gov.au/Infectious/controlguideline/Pages/hendra-case-summary.aspx

8. Arunkumar G, Chandni R, Mourya DT, Singh SK, Sadanandan R, Sudan P, et al. Outbreak Investigation of Nipah Virus Disease in Kerala, India, 2018. J Infect Dis. 2019 May 24;219(12):1867–78.

9. Ching PK, de los Reyes VC, Sucaldito MN, Tayag E, Columna-Vingno AB, et al. 2015. Outbreak of henipavirus infection, Philippines, 2014. Emerg. Infect. Dis. 21:328–31

10. Nikolay B, Salje H, Hossain MJ, Khan AKMD, Sazzad HMS, Rahman M, et al. Transmission of Nipah Virus — 14 Years of Investigations in Bangladesh. New England Journal of Medicine. 2019 May 9;380(19):1804–14.

11. Amaya M, Broder CC. Vaccines to Emerging Viruses: Nipah and Hendra. Annu Rev Virol. 2020 Sep 29;7(1):447–73.

12. Middleton D, Pallister J, Klein R, Feng YR, Haining J, Arkinstall R, et al. Hendra virus vaccine, a one health approach to protecting horse, human, and environmental health. Emerging Infectious Diseases. 2014 Mar;20(3):372–9.

13. Playford EG, Munro T, Mahler SM, Elliott S, Gerometta M, Hoger KL, et al. Safety, tolerability, pharmacokinetics, and immunogenicity of a human monoclonal antibody targeting the G glycoprotein of henipaviruses in healthy adults: a first-in-human, randomised, controlled, phase 1 study. The Lancet Infectious Diseases. 2020;20(4):445–54.

14. Dong J, Cross RW, Doyle MP, Kose N, Mousa JJ, Annand EJ, et al. Potent henipavirus neutralization by antibodies recognizing diverse sites on Hendra and Nipah virus attachment glycoproteins. Cell. 2020;183:1536–50.

15. Doyle MP, Kose N, Borisevich V, Binshtein E, Amaya M, Nagel M, et al. Functional cooperativity mediated by rationally selected combinations of human monoclonal antibodies targeting the henipavirus receptor binding protein. Cell Reports. 2021 Aug 31;36(9):109628.

16. Dang HV, Cross RW, Borisevich V, Bornholdt ZA, West BR, Chan Y-P, et al. Broadly neutralizing antibody cocktails targeting Nipah virus and Hendra virus fusion glycoproteins. Nature Structural & Molecular Biology. 2021;28(5):426–34.

17. Geisbert TW, Bobb K, Borisevich V, Geisbert JB, Agans KN, Cross RW, et al. A single dose investigational subunit vaccine for human use against Nipah virus and Hendra virus. npj Vaccines. 2021 Dec;6(1):23

18. Kirkland PD, Gabor M, Poe I, Neale K, Chaffey K, Finlaison DS, et al. Hendra Virus Infection in Dog, Australia, 2013. Emerg Infect Dis. 2015 Dec;21(12):2182–5.

19. Edson D, Field H, McMichael L, Vidgen M, Goldspink L, Broos A, et al. Routes of Hendra virus excretion in naturally-infected flying-foxes: implications for viral transmission and spillover risk. PLoS One [Internet]. 2015 Oct 15 [cited 2021 May 21];10(10):e0140670. Available from: https://www.ncbi.nlm.nih.gov/pmc/articles/PMC4607162/

20. Burroughs AL, Durr PA, Boyd V, Graham K, White JR, Todd S, et al. Hendra Virus infection dynamics in the grey-headed flying fox (Pteropus poliocephalus) at the Southern-most extent of its range: further evidence this species does not readily transmit the virus to horses. PLoS One. 11(6):e0155252.

21. Edson D, Peel AJ, Huth L, Mayer DG, Vidgen ME, McMichael L, et al. Time of year, age class and body condition predict Hendra virus infection in Australian black flying foxes (Pteropus alecto). Epidemiology and Infection [Internet]. 2019 Jul 10;147:e240. [cited 2021 May 28] Available from: https://www.ncbi.nlm.nih.gov/pmc/articles/PMC6625375/

22. Boardman WSJ, Baker ML, Boyd V, Crameri G, Peck GR, Reardon T, et al. Seroprevalence of three paramyxoviruses; Hendra virus, Tioman virus, Cedar virus and a rhabdovirus, Australian bat lyssavirus, in a range expanding fruit bat, the Grey-headed flying fox (*Pteropus poliocephalus*). PLoS One. 2020;15(5):e0232339.

23. Plowright RK, Eby P, Hudson PJ, Smith IL, Westcott D, Bryden WL, et al. Ecological dynamics of emerging bat virus spillover. Proceedings of the Royal Society B. 2015;282.

24. Annand EJ, Reid PA, Johnson J, Gilbert GL, Taylor M, Walsh M, et al. Citizens’ juries give verdict on whether private practice veterinarians should attend unvaccinated Hendra virus suspect horses. Australian Veterinary Journal. 2020;98(7):273–9.

25. Animal Health Australia. Animal Health Surveillance Quarterly – Animal Health Australia [Internet]. [cited 2021 May 21]. Available from: https://www.animalhealthaustralia.com.au/our-publications/animal-health-surveillance-quarterly/

26. Government of South Australia. Department of Primary Industries and Regions. Hendra virus in South Australia [Internet]. 2018 [cited 2021 May 21]. Available from: https://pir.sa.gov.au/biosecurity/animal_health/horses/hendra_virus

27. Agnihotri K, Pease B, Oakey J, Campbell G. Confirmation of Elsey virus infection in a Queensland horse with mild neurological signs. Journal of Veterinary Diagnostic Investigation. 2016;28(4):445–8.

28. Smith IL, Halpin K, Warrilow D, Smith GA. Development of a fluorogenic RT-PCR assay (TaqMan) for the detection of Hendra virus. Journal of Virological Methods. 2001;98(1):33–40.

29. Feldman KS, Foord A, Heine HG, Smith IL, Boyd V, Marsh GA, et al. Design and evaluation of consensus PCR assays for henipaviruses. Journal of Virological Methods. 2009 Oct 1;161(1):52–7.

30. Yuen KY, Fraser NS, Henning J, Halpin K, Gibson JS, Betzien L, et al. Hendra virus: epidemiology dynamics in relation to climate change, diagnostic tests and control measures. One Health [Internet]. 2020 Dec 21 [cited 2021 May 21];12. Available from: https://www.ncbi.nlm.nih.gov/pmc/articles/PMC7750128/

31. Annand EA, Reid PA. Clinical review of two fatal equine cases of infection with the insectivorous bat strain of Australian bat lyssavirus. Australian Veterinary Journal. 2014;92(9):324–32.

32. Annand E, Barr J, Singanallur Balasubramanian N, Reid P, Boyd V, Burneikienė-Petraitytė R, et al. Spillover of bat borne Rubulavirus in Australian horses – Horses as sentinels for emerging infectious diseases. International Journal of Infectious Diseases. 2020 Dec 1;101:401–2.

33. Barr J, Smith C, Smith I, de Jong C, Todd S, Melville D, et al. Isolation of multiple novel paramyxoviruses from pteropid bat urine. Journal of General Virology. 2015;96(Pt 1):24–9.

34. Marsh GA, Jong CD, Barr J, Tachedjian M, Smith C, Middleton D, et al. Cedar virus: a novel henipavirus isolated from Australian bats. PLOS Pathogens. 2012;8(8):e1002836.

35. Vidgen ME, Jong CD, Rose K, Hall J, Field H, Smith C. Novel paramyxoviruses in Australian flying-fox populations support host–virus coevolution. Journal of General Virology. 2015;96:1619–25.

36. Tong S, Chern S-WW, Li Y, Pallansch MA, Anderson LJ. Sensitive and broadly reactive reverse transcription-PCR assays to detect novel paramyxoviruses. Journal of Clinical Microbiology. 2008;46(8):2652–8.

37. Li D, Liu C-M, Luo R, Sadakane K, Lam T-W. MEGAHIT: an ultra-fast single-node solution for large and complex metagenomics assembly via succinct de Bruijn graph. Bioinformatics. 2015;31(10):1674–6.

38. Altschul SF, Gish W, Miller W, Myers EW, Lipman DJ. Basic local alignment search tool. Journal of Molecular Biology. 1990;215(3):403–10.

39. Rima B, Balkema-Buschmann A, Dundon WG, Duprex P, Easton A, Fouchier R, et al. ICTV virus taxonomy profile: paramyxoviridae. Journal of General Virology. 2019;100(12):1593–4.

40. Crameri G, Todd S, Grimley S, McEachern JA, Marsh GA, Smith C, et al. Establishment, immortalisation and characterisation of pteropid bat cell lines. PLoS One. 2009;4(12):e8266.

41. Xu K, Rockx B, Xie Y, DeBuysscher BL, Fusco DL, Zhu Z, et al. Crystal structure of the Hendra virus attachment G glycoprotein bound to a potent cross-reactive neutralizing human monoclonal antibody. PLoS Pathogens [Internet]. 2013 Oct 10 [cited 2021 Apr 27];9(10). Available from: https://www.ncbi.nlm.nih.gov/pmc/articles/PMC3795035/

42. Australian Government Department of Agriculture, Water and the Environment. National Flying-Fox Monitoring Program [Internet]. Monitoring Flying-Fox Populations. [cited 2021 May 28]. Available from: https://www.environment.gov.au/biodiversity/threatened/species/flying-fox-monitoring

43. Colling A, Lunt R, Bergfeld J, McNabb L, Halpin K, Juzva S, et al. A network approach for provisional assay recognition of a Hendra virus antibody ELISA: test validation with low sample numbers from infected horses. Journal of Veterinary Diagnostic Investigation. 2018;30(3):362–9.

44. Hyatt AD, Zaki SR, Goldsmith CS, Wise TG, Hengstberger SG. Ultrastructure of Hendra virus and Nipah virus within cultured cells and host animals. Microbes Infect. 2001 Apr;3(4):297–306. doi: 10.1016/s1286-4579(01)01383-1. PMID: 11334747.

45. Laing E, Yan L, Sterling S, Broder C. A Luminex-based multiplex assay for the simultaneous detection of glycoprotein specific antibodies to ebolaviruses, marburgviruses, and henipaviruses. International Journal of Infectious Diseases. 2016 Dec 1;53:108–9.

46. Wang J, Anderson D, Valdeter S, Chen H, Walker S, Meehan B, et al. A novel henipavirus in bats, Australia. In: Proceedings of the One Health EcoHealth Congress [Internet]. Melbourne: One Health EcoHealth, 2016 [cited 2021 Feb 18]. Available from: https://publications.csiro.au/rpr/pub?pid=csiro:EP173003

47. Wang J, Anderson DE, Halpin K, Hong X, Chen H, Walker S, et al. A New Hendra Virus Genotype Found in Australian Flying Foxes. Virology Journal, In Review. Research Square Pre-print DOI:10.21203/rs.3.rs-616496.

48. Roberts BJ, Catterall CP, Eby P, Kanowski J. Latitudinal range shifts in Australian flying-foxes: a re-evaluation. Austral Ecology. 2012;37(1):12–22.

49. Mendez DH, Judd J, Speare R. Unexpected result of Hendra virus outbreaks for veterinarians, Queensland, Australia. Emerging Infectious Diseases. 2012;18(1):83–5.

50. Hendra Virus Interagency Technical Working Group. Hendra Virus Infection Prevention Advice - Hendra Virus Interagency Technical Working Group incl. Biosecurity Queensland, Australian Veterinary Association, Queensland Health, Workplace Health & Safety Queensland [Internet]. Queensland Government; 2014. Available from: https://www.health.qld.gov.au/_data/assets/pdf_file/0026/428624/hev-inf-prev-adv.pdf

51. Animal Health Australia. Australia Veterinary Emergency Plan AUSVETPLAN, Response policy brief Hendra virus infection Version 4.0. [Internet]. Animal Health Australia; 2016 [cited 2021 May 5]. Available from: https://www.animalhealthaustralia.com.au/download/2602/

